# Structure of human Na_v_1.5 reveals the fast inactivation-related segments as a mutational hotspot for the Long QT Syndrome

**DOI:** 10.1101/2021.02.06.430010

**Authors:** Zhangqiang Li, Xueqin Jin, Tong Wu, Xin Zhao, Weipeng Wang, Jianlin Lei, Xiaojing Pan, Nieng Yan

## Abstract

Na_v_1.5 is the primary voltage-gated Na^+^ (Na_v_) channel in the heart. Mutations of Na_v_1.5 are associated with various cardiac disorders exemplified by the type 3 long QT syndrome (LQT3) and Brugada syndrome (BrS). E1784K is a common mutation that has been found in both LQT3 and BrS patients. Here we present the cryo-EM structure of the human Na_v_1.5-E1784K variant at an overall resolution of 3.3 Å. Structural mapping of 91 and 178 point mutations that are respectively associated with LQT3 and BrS reveals a unique distribution pattern for LQT3 mutations. Whereas the BrS mutations spread evenly on the structure, LQT3 mutations are mainly clustered to the segments in repeats III and IV that are involved in gating, voltage-sensing, and particularly inactivation. A mutational hotspot involving the fast inactivation segments is identified and can be mechanistically interpreted by our “door wedge” model for fast inactivation. The structural analysis presented here, with a focus on the impact of disease mutations on inactivation and late sodium current, establishes a structure-function relationship for the mechanistic understanding of Na_v_1.5 channelopathies.

## Introduction

Cardiac arrhythmia affects 1-2% world populations and accounts for 230,000-350,000 sudden death in the United States each year (1, 2). A leading cause for arrythmia is aberrant firing of electrical signals, mediated by the voltage-gated Na channel Na_v_1.5 encoded by *SCN5A*. Dysfunction of Na_v_1.5 is related to a variety of arrhythmia syndromes, such as the Brugada syndrome (BrS), the long QT syndrome (LQTS), and the Sick Sinus syndrome (SSS) (3-5) (*SI Appendix*, Tables S1-S3). Congenital LQTS and BrS are the two most prevalent genetic disorders for cardiac channelopathies that account for approximately half of sudden arrhythmic death incidents (6-8).

LQT was named for the phenotype of prolonged Q-T intervals on the electrocardiography (ECG) that was first reported more than six decades ago (9). Ensuing studies revealed that functional alteration of channels that control repolarization underlay many hereditary LQT cases. LQT3, with arrhythmia during sleep or upon waking, represents the most harmful LQT type. Inherited LQT3 is mainly associated with mutations in Na_v_1.5, many of which result in increased late Na^+^ current (*I*_*Na*_), a form of gain of function (GOF) (10, 11). In contrast, BrS-associated Na_v_1.5 mutations are mostly loss of function (LOF), manifested by reduced peak *I*_*Na*_ or leftward shift of inactivation curves (12-14).

Since the identification of the LQTS-related *SCN5A* mutation in 1995, more than 400 missense mutations in the primary sequence of Na_v_1.5 have been reported related to various cardiac disorders (10, 15). Among these, 154 and 230 mutations are respectively identified in patients with LQT3 and BrS (*SI Appendix*, Tables S1, S2). Intriguingly, approximately 30 mutations were found in both LQT3 and BrS (*SI appendix*, Tables S1, S2).

To further the molecular understanding of Na_v_1.5 channelopathies, we sought to solve structures of human Na_v_1.5 with disease mutations in addition to our previous structural resolution of wild type (WT) human Na_v_1.5 bound to pore blockers (16). We started with Na_v_1.5-E1784K, which is the most common mutation shared by LQT3 and BrS. This mutation enhances the risk of sudden death resulted from ventricular tachyarrhythmias in LQT3 patients or idiopathic ventricular fibrillation in BrS patients (17, 18). Recombinantly expressed Na_v_1.5-E1784K exhibited leftward shift (∼15 mV) of the steady-state inactivation, reduced peak *I*_*Na*_, and increased late *I*_*Na*_ (17, 19, 20). How one single point mutation leads to both GOF and LOF at the same time remains enigmatic.

In this paper, we report the structural determination of Na_v_1.5-E1784K using single particle cryo-electron microscopy (cryo-EM). Mapping of disease mutations shows distinct distribution patterns for LQT3 and BrS mutations on the three dimensional structure. The fast inactivation-related structural elements stand out as a hsotspot for LQT3 mutations.

## Results

### Cryo-EM Analysis of Human Na_v_1.5-E1784K

Consistent with previous report, substitution of Glu1784 with Lys led to a leftward shift (∼20 mV) of the steady-state inactivation, increased late *I*_*Na*_ at room temperature, and significantly decreased conductance of Na_v_1.5 compared to the WT channel (17, 19, 20) (Fig 1*A**, SI Appendix*, Fig S1 and Table S4).

**Figure 1.**
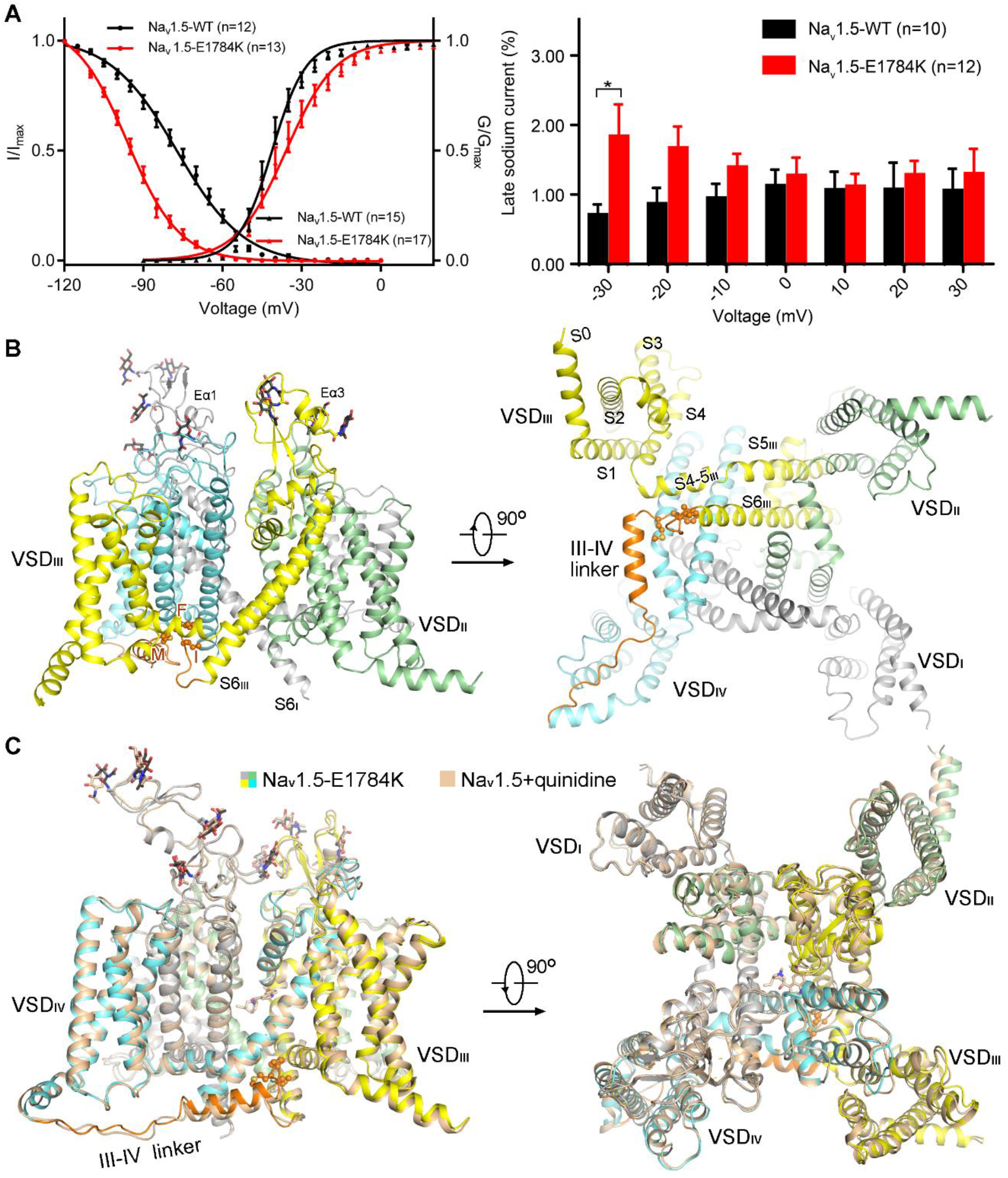
Structure of full-length human Na_v_1.5-E1784K. (*A*) Electrophysiological properties of Na_v_1.5-E1784K heterogeneously expressed in HEK293T cells. *left*: Voltage-dependent activation and inactivation curves. Detailed parameters are presented in *SI appendix*, Table S4. *right*: Persistent sodium current at different voltages. The working temperature is 22 °C. (*B*) Overall structure of human Na_v_1.5-E1784K. A side and a cytoplasmic view are shown. The structure is color-coded for distinct repeats. The III-IV linker is colored orange and the fast inactivation motif, Ile/Phe/Met (IFM) is shown as ball and sticks on the right. The sugar moieties and is shown as black and blue sticks, respectively. Eα1/3: Extracellular α helix in repeat I or III. All structural figures are prepared in PyMol (57). (*C*) The structure of Na_v_1.5-E1784K is nearly identical with that of Na_v_1.5-qunidine (PDB code: 6LQA). The complex structures of Na_v_1.5-E1784K and Na_v_1.5-qunidine can be superimposed with an RMSD of 0.708 Å over 1075 Cα atoms.

Four auxiliary β subunits, β1-β4, modulate the localization and channel properties of the α core subunit of Na_v_ channels (21). We co-expressed β1 with Na_v_1.5-E1784K to improve the expression level, although there was no corresponding density for any auxiliary subunit in the 3D EM map. Details for protein preparation, image acquisition, and data processing are presented in Methods. A 3D EM reconstruction was attained at the overall resolution of 3.3 Å out of 147,600 selected particles (*SI Appendix*, Figs S2-S3 and Table S5).

The final structural model of Na_v_1.5-E1784K contains 1151 residues, including the complete transmembrane domain, the extracellular segments, and the III-IV linker. Nine glycosylation sites were observed and assigned to each site (Fig. 1*B*). Despite the change of the channel properties, the structure of Na_v_1.5-E1784K is nearly identical to that of quinidine-bound WT Na_v_1.5 (Fig. 1*C*) (16). More importantly, the last resolved residue is Glu1781 that marks the end of S6_IV_. The invisibility of Glu(Lys)1784 prevents a direct structural interpretation of the pathogenic mechanism of this mutation. We therefore shift our focus to systematic structure-based analysis of resolved mutation sites, in the hope to facilitate the establishment of a structure-function relationship of Na_v_1.5 disease variants.

### Distinct Structural Distributions of LQT3 and BrS Mutations

The high-resolution structure of human Na_v_1.5-E1784K allows for mapping of a total of 86 and 174 mutations that are associated with LQT3 and BrS, respectively (*SI Appendix*, Tables S1-S2).

Structural mapping of the disease mutations reveals distinctive distribution patterns of BrS and LQT3 mutations. The BrS mutations span the entire structure, including 39 mutations affecting 32 residues on the extracellular loops (ECL) and 30 mutations of 24 residues on the P1-SF-P2 region (described as the SF zone hereafter) (Fig. 2*A*, *SI Appendix*, Tables S1-S2). In contrast, there are only 6 and 3 LQT3 mutations identified on the ECL and SF region, respectively (Fig. 2*B*). In fact, the structural distribution of LQT3 mutations appears to be highly polarized, with 33 residues (37 mutations) clustered on the III-IV linker and the nearby S4-S5, S5, and S6 segments in repeats III and IV (Fig. 3*B* and *C*). In the following session, we will focus on a structural overview of LQT3 mutations.

**Figure 2.**
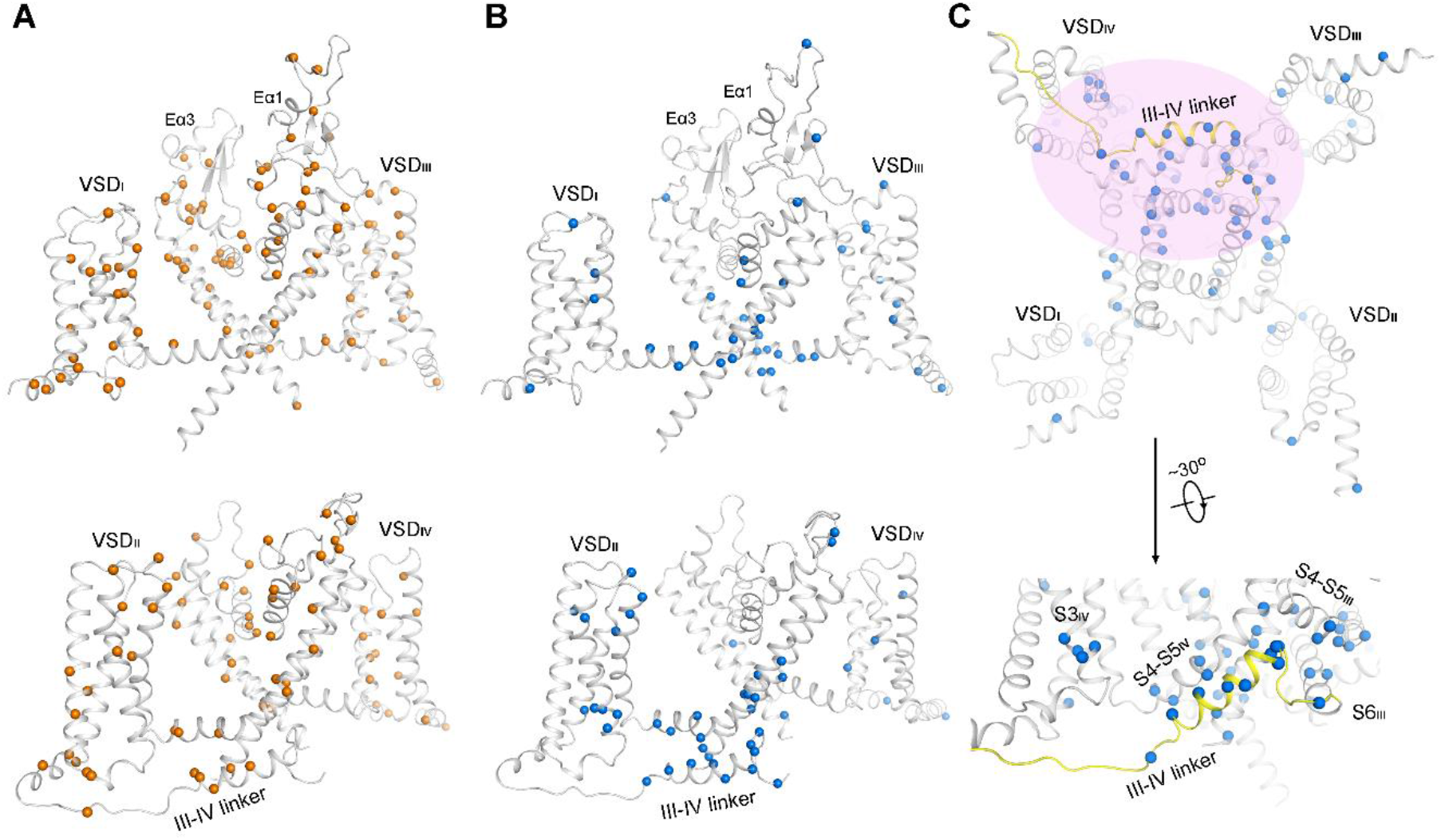
Structural mapping of Na_v_1.5 mutations that are associated with Brugada syndrome (BrS) and LQT3. (*A* and *B*) Mapping of the BrS (*A*) and LQT3 (*B*) disease mutations on the structure. The Cα atoms of the disease-related residues are shown as spheres. Diagonal repeats are shown on the top and bottom rows. Whereas BrS mutations are distributed throughout the structure, LQT3 mutations are mainly found in the VSDs and the segments related to fast inactivation. (*C*) The Ile/Phe/Met (IFM) motif-carrying III-IV linker and its receptor site represents a hotspot for LQT3 mutations. The cluster of mutations is indicated by the semi-transparent pink oval. The III-IV linker is colored yellow.

**Figure 3.**
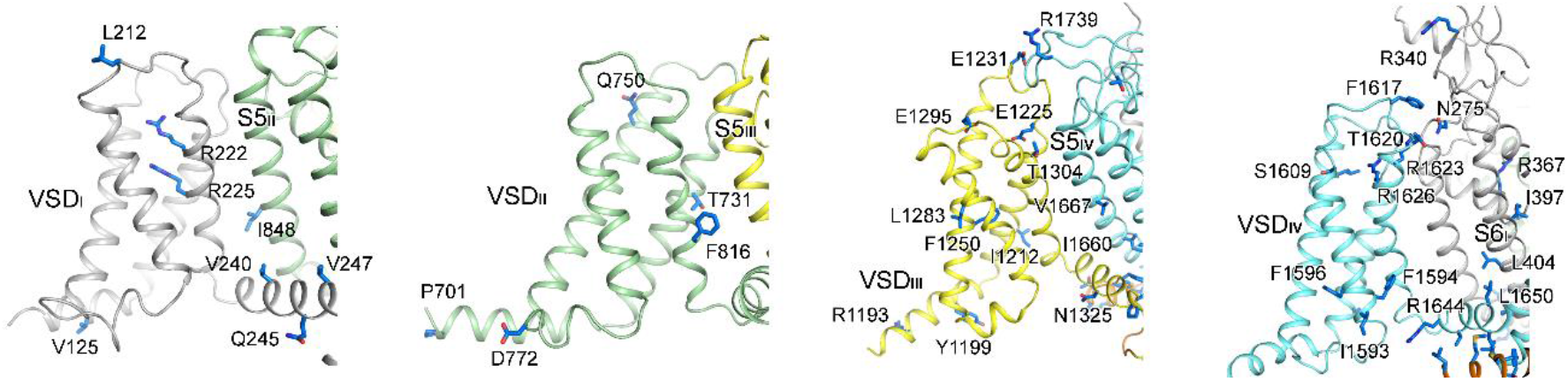
Mapping of LQT3 mutations on VSDs. VSD_III_ and VSD_IV_ harbor more LQT3 mutations than the other two VSDs. The LQT3-related residues that are mapped to VSDs, including the S4-S5 linker, and their respective interface with the PD are shown as blue sticks.

### LQT3 Mutations in voltage-sensing domains (VSDs)

An asynchronous activation model of the four VSDs has been established for Na_v_ channel, whereby the first three VSDs activate in advance of VSD_IV_. Whereas activation of the first three VSDs leads to pore opening, that of VSD_IV_ elicits fast inactivation of the channel (22-25). Consistent with their distinct functions, distribution of LQT3-related residues is uneven among the four VSDs, with four on VSD_I_, five on VSD_II_, nine on VSD_III_, and ten on VSD_IV_ (Fig. 3, *SI Appendix*, Table S1). Of note, counted here are the affected loci, although multiple substitutions can occur to the same residue. None of the mutations in VSD_II_ and VSD_III_ involves the well-defined functional residues such as the gating charge (GC) residues, whereas two of the four affected residues are GC residues R2 and R3 (R222Q, R225Q/W) in VSD_I_ (26, 27). Similarly, mutations of two GCs, R1 and R2, in VSD_IV_ are found in LQT3 patients (R1623L/Q, R1626H/P) (11, 28, 29) (*SI Appendix*, Table S1).

Mechanistic dissection of the LQT3 mutations in VSDs, which undergo pronounced conformational changes during the working cycle of Na_v_ channels for each firing, requires structural determination of the channel in multiple functioning states. Prior to that, comparison of the structures of Na_v_1.5-E1784K and Na_v_PaS, which exhibits a conformation that is distinct from all mammalian Na_v_ channels of known structures, affords important clue to understanding the primary mutational hotspot on Na_v_1.5 for LQT3.

### A LQT3 Mutational Hotspot Mapped to the Inactivation Segments

An important discovery derived from our comprehensive structural mapping of disease mutations is that approximately 20% of all LQT3 mutations, or about one third of structurally resolved mutations, are clustered to one corner of the pore domain (PD), which is close to the intracellular gate along the permeation path in repeats III and IV (Fig. 4*A*). In particular, the Ile/Phe/Met (IFM) motif and the ensuing α helix on the III-IV linker host an exceptionally high density of LQT3 mutations (Fig. 4*B*).

**Figure 4.**
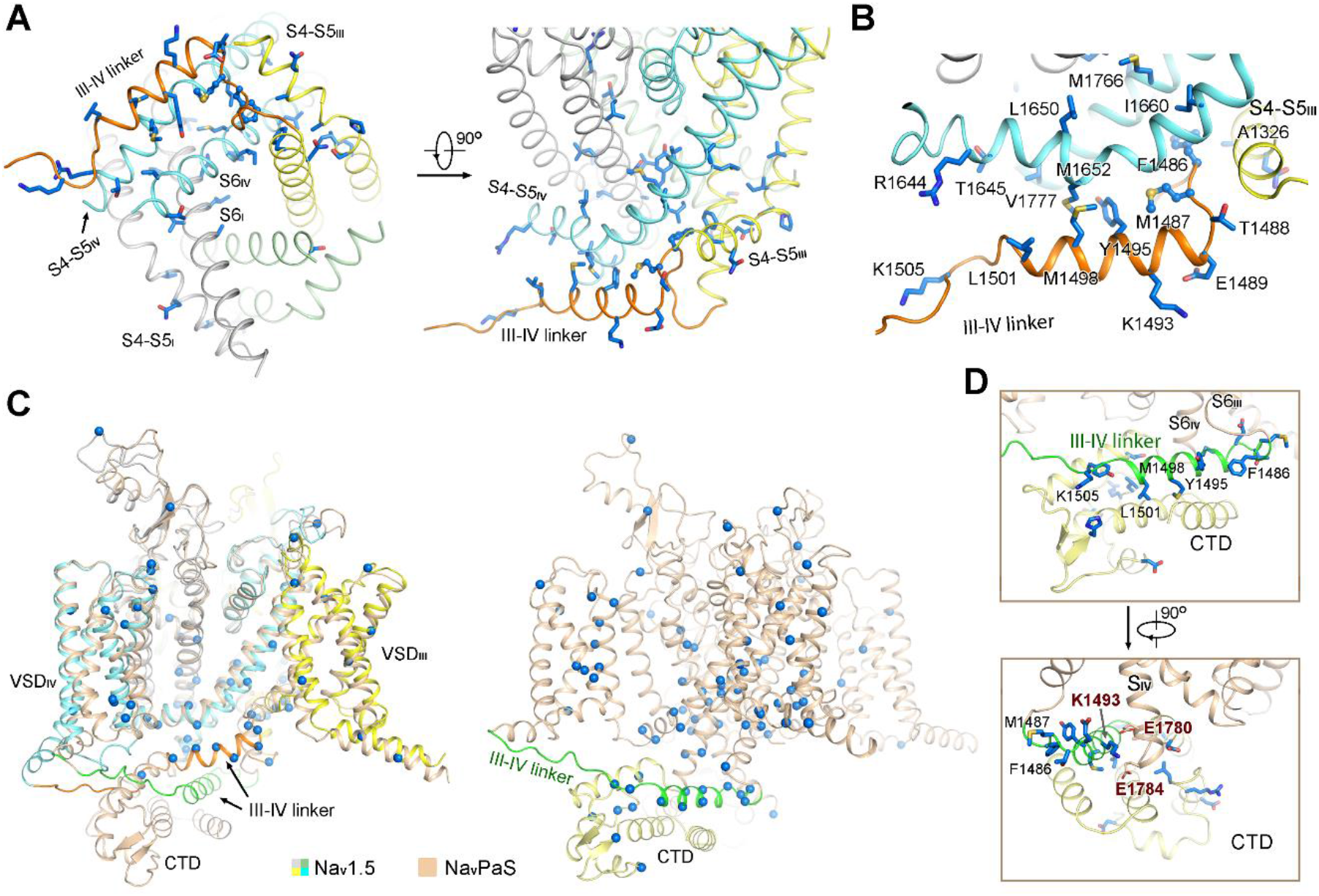
Structural analysis of fast inactivation-related LQT3 mutations. (*A*) Asymmetric distribution of LQT3 mutations on the pore domain. The LQT3-related residues on the pore domain are shown as blue sticks. An intracellular (left) and a side (right) view are presented. These mutations are clustered to the gating and fast inactivation-related segments in repeats III and IV. (*B*) The interface of the III-IV linker and S4-S5_IV_ is enriched of LQT3 mutations. (*C*) Mapping of the Na_v_1.5 LQT3 mutations to Na_v_PaS. The structure of Na_v_PaS, a Na_v_ channel from American cockroach, exhibits a distinct conformation from that of Na_v_1.5, featured with less depolarized VSDs, sealed PD, and the C-terminal domain (CTD)-sequestered III-IV linker (PDB code: 5X0M). *Left*: Structural superimposition of Na_v_1.5-E1784K and Na_v_PaS. *Right*: Many LQT3-related residues may be mapped to the interface of the III-IV linker and the CTD. The Cα atoms of the Na_v_PaS residues that correspond to the LQT3 mutations in Na_v_1.5 are shown as spheres. (*D*) Mapping of LQT3 mutations in a Na_v_PaS-derived Na_v_1.5 model, in which the conformation of the PD and the III-IV linker may represent a potentially resting state. *Upper*: A Na_v_PaS-derived structural model for Na_v_1.5. The LQT3-related residues that map to the interface of the III-IV linker and the CTD are shown as blue sticks. *Lower*: Lys1493 on the III-IV helix may be sandwiched by the invariant Glu1780 and Glu1784 on the end of S6_IV_ in a potentially resting state. Mutation K1493R may strengthen these interactions, hence impeding the conformational shift towards the inactivation conformation captured by the structure, as seen in panel (*B*). Consistently, the LQT3 mutation K1493R led to right shift of the steady-state inactivation curve (41). Glu1784 may also be involved in the interaction with CTD and the III-IV linker. Mutation E1784K may disrupt this interaction, resulting in accelerated fast inactivation and had a leftward shift of the inactivation curve, but the molecular basis for the increased late *I*_*Na*_ awaits further characterizations (Fig. 1*A*) (17, 42).

Decades of research have identified that III-IV linker plays a fundamental role for fast inactivation of Na_v_ channels, among which the hydrophobic cluster IFM motif was defined as the fast inactivation particle (30, 31). In contrast to the conventional “ball-and-chain” model (31), our structural comparison of Na_v_PaS with other Na_v_ channels suggested an “allosteric blocking” mechanism (32, 33), or “door wedge” model, for fast inactivating Na_v_ channels. In Na_v_PaS, the III-IV linker is sequestered by the carboxy terminal domain (CTD) (33, 34), leaving the IFM-corresponding residues away from the PD, a conformation that is also observed in the voltage-gated Ca^2+^ channel Ca_v_1.1 (35) (Fig. 4*C*). In the structures of EeNa_v_1.4 and all resolved human Na_v_ channels (16, 32, 36-38), the IFM motif undergoes a pronounced displacement to plug into a cavity that is constituted by the S6 helices and the S4-S5 segments in repeats III and IV (Fig. 4*C*). Insertion of the IFM motif into this receptor site outside the intracellular gate is expected to push the S6 segments to close at the intracellular gate (33), reminiscent of a door stopper wedge. Therefore, these structures, characteristic with “up” VSDs and plugged IFM motif, may all represent the inactivated state of Na_v_ channels. For simplicity, we will refer to the IFM motif and its receptor site as the “fast inactivation corner”.

Based on our “door wedge” model, we have the following hypothesis: mutations that impair interactions between the IFM motif-carrying III-IV linker and its receptor site, relax the S4-S5 restriction ring surrounding the intracellular gate, or disrupt the intracellular gate may hinder fast inactivation and enhance the probability of gate opening during prolonged depolarization. These molecular events would be detected, in the electrophysiological characterizations, as right shift of the steady-state inactivation curve and increased late *I*_*Na*_, which are just the commonly observed channel alterations associated with LQT3 mutations.

Our analysis is supported by documented characterizations of Na_v_1.5 variants containing LQT3 mutations. For instance, mutation of the Phe in the IFM motif, F1486L, or alternation of a Phe in its receptor cavity, F1473C, may decrease the affinity between IFM and the receptor site. Both mutations led to increased late *I*_*Na*_ (39, 40). Met1652 on the S4-S5_IV_ segment interacts with Tyr1495 and Met1498 on the III-IV linker (Fig. 4*B*). Single point mutation M1652R, which may disrupt the interaction with the III-IV linker, resulted in slowed inactivation and increased late *I*_*Na*_ (11).

Along the same line, mutations that stabilize the resting state and impede conformational shifts toward the inactivated structure may also lead to LQT3 phenotype. Before the structure of any eukaryotic Na_v_ channel in the resting-state is captured, that of Na_v_PaS structure affords important clues (Fig. 4*C* and *D*) (33, 34). To facilitate structural mapping, a 3D model of Na_v_1.5 is generated based on sequence homology with Na_v_PaS (PDB code: 5X0M).

In this Na_v_PaS-derived Na_v_1.5 model, the III-IV linker interacts with the CTD. Although several LQT3 mutations are mapped to the CTD (Fig. 4*D*, upper), we will refrain from discussing them because the resolution of CTD did not support accurate side chain modelling in Na_v_PaS (33, 34). We focus on the residues whose counterparts in Na_v_PaS were reliably resolved. Na_v_1.5-Lys1493, which is on the III-IV linker helix and away from the interface with other structural segment in the cryo-EM structure of Na_v_1.5-E1784K (Fig. 4*B*), would be sandwiched by Glu1780 and Glu1784, which are on the C-terminal end of S6_IV_, in the conformation of Na_v_PaS (Fig. 4*D*, lower). Replacement of Lys1493 by Arg may even strengthen these interactions, hence impeding the dislocation of the III-IV linker toward an inactivated conformation. Indeed, the LQT3 mutation K1493R resulted in a right shift of the steady-state inactivation curve (41).

Na_v_1.5-Glu1784, albeit invisible in the current structure, is predicted to be positioned at the cytosolic tip of S6_IV_. Based on the Na_v_PaS-derived model, replacement of the acidic residue with Lys may weaken the sequestration of the S6_IV_ segment by the CTD, resulting in accelerated fast inactivation and left shift of the inactivation curve, which may account for the BrS phenotype. However, the overall impact of this single point mutation may be more complex, because it also led to increased late *I*_*Na*_ underlying the LQT3 phenotype that awaits a mechanistic explanation (17, 42).

## Discussion

In this paper, we report the structure of human Na_v_1.5-E1784K and focus on the systematic structural analysis of LQT3-associated mutations. Instead of characterizing specific disease mutations, we attempted to summarize the distributing pattern of the resolved LQT3 mutations to acquire mechanistic insight. The structural analysis, together with reported functional characterizations, helps establish a structure-function relationship of the pathogenic mutations and in turn illuminates the mechanistic understanding of the Na_v_ channels.

An advanced molecular dissection of these pathogenic mutations necessitates structural determination of Na_v_ channels in multiple functional states. Among all the structures of eukaryotic Na_v_ channels, only two major conformations were captured. The structure of Na_v_PaS is featured with closed PD without fenestration, a CTD-sequestered III-IV linker, and VSDs in less polarized conformations (33, 34); while all the human, electric eel, and rat Na_v_ channels exhibit similar conformations characterized by the insertion of the IFM motif into a cavity outside the S6 tetrahelical bundle (16, 32, 36-38, 43). Although the functional state of the Na_v_PaS structure cannot be defined due to its resistance to electrophysiological recording, the conformations of the PD, CTD, and the III-IV linker are consistent with those expected in a resting or pre-open channel. The pronounced structural differences between Na_v_PaS and other Na_v_ channels thus provide the opportunity for understanding the function and disease mechanism of Na_v_ channels.

Structural mapping reveals the fast inactivation-related segments to be a hotspot for LQT3 mutations. Our “door wedge” model for fast inactivation, originally derived from structural comparison of Na_v_Pas and Na_v_1.4 channels (32, 37) and supported by functional data reported in the past three decades, affords a plausible mechanistic interpretation for the exceptionally high-density of LQT3 mutations on the III-IV linker and the surrounding segments.

## Materials and Methods

### Transient Co-expression of Human Na_v_1.5-E1784K and β1

A single point mutation E1784K was introduced to full-length human Na_v_1.5 (Uniprot: Q14524) by QuickChange site-directed mutagenesis and confirmed by PCR sequencing, then the cDNAs was cloned into the pEG BacMam vector with twin Strep-tag and FLAG tag in tandem at the amino terminus, and β1 (Uniprot: Q07699) was cloned into the pCAG vector without any affinity tag (44, 45). Both the cDNAs of Na_v_1.5-E1784K and β1 were optimized into mammalian cell expression system and co-expressed in HEK293F cells (Invitrogen), which were cultured in SMM 293T-II medium (Sino Biological Inc.) under 5% CO_2_ in a Multitron-Pro shaker (Infors, 130 r.p.m.) at 37 °C. When the cell density reached about 2.0 × 10^6^ cells per mL, approximately 2.0 mg plasmids (1.4 mg for Na_v_1.5-E1784K and 0.6 mg for β1) and 4 mg 25-kDa linear polyethylenimines (PEIs) (Polysciences) were mixed into 30 mL fresh medium and pre-incubated for 15-30 min before adding into 1 liter cell culture. Transfected cells were harvested after 48 h cultivation.

### Protein Purification of Na_v_1.5-E1784K and β1

Protein purification, sample preparation, data collection, and model building was conducted following a protocol identical to that for WT human Na_v_1.5 channel (16), which was derived from the experiments for human Na_v_1.4 (37). 24 L transfected cells were harvested by centrifugation at 800 g for 12 min and the cell pellet was resuspended in the lysis buffer containing 25 mM Tris-HCl (pH 7.5) and 150 mM NaCl. The cell suspension was supplemented with 1% (w/v) n-dodecyl-β-D-maltopyranoside (DDM, Anatrace), 0.1% (w/v) cholesteryl hemisuccinate Tris salt (CHS, Anatrace), and protease inhibitor cocktail containing 2 mM phenylmethylsulfonyl fluoride (PMSF), aprotinin (3.9 μg/mL), pepstatin (2.1 μg/mL), and leupeptin (15 μg/mL), and incubated at 4 °C for 3 h. The cell lysate was subsequently ultra-centrifuged at 20,000 g for 45 min, and the supernatant was applied to anti-Flag M2 affinity gel (Sigma) by gravity at 4 °C. The resin was washed with 10 column volumes of wash buffer containing 25 mM Tris-HCl (pH 7.5), 150 mM NaCl, 0.06% glycol-diosgenin (GDN, Anatrace), and the protease inhibitor cocktail. Target proteins were eluted with 5 column volumes of wash buffer plus 200 μg/mL FLAG peptide (Sigma). The eluent was then applied to Strep-Tactin Sepharose (IBA) and the purification protocol was much similar with the previous steps except the elution buffer, which was wash buffer plus 2.5 mM D-Desthiobiotin (IBA). The eluent was then concentrated using a 100-kDa cut-off Centricon (Millipore) and further purified with SEC (Superose-6 Increase 10/300 column, GE Healthcare) in the wash buffer. The presence of target proteins was confirmed by SDS-PAGE and mass spectrometry. The purified proteins were pooled and concentrated to approximately 2 mg/mL for cryo-EM sample preparation.

### Whole Cell Electrophysiology

HEK293T cells (Invitrogen) cultured in Dulbecco’s Modified Eagle Medium (DMEM, BI) supplemented with 4.5 mg/mL glucose and 10% fetal bovine serum (FBS, BI) were plated onto glass coverslips for subsequent patch clamp recordings. Cells were transiently co-transfected using lipofectamine 2000 (Invitrogen) with the expression plasmids for indicated Na_v_1.5 or Na_v_1.5-E1784K and an eGFP-encoding plasmid. Cells with green fluorescence were selected for patch-clamp recording at 18–36 h after transfection. All experiments were performed at room temperature. No further authentication was performed for the commercially available cell line. Mycoplasma contamination was not tested.

The whole-cell Na^+^ currents were recorded in HEK293T cells using an EPC10-USB amplifier with Patchmaster software v2*90.2 (HEKA Elektronic), filtered at 3 kHz (low-pass Bessel filter) and sampled at 50 kHz. The borosilicate pipettes (Sutter Instrument) used had a resistance of 2-4 MΩ and the electrodes were filled with the internal solution composed of (in mM) 105 CsF, 40 CsCl, 10 NaCl, 10 EGTA, 10 HEPES, pH 7.4 with CsOH. The bath solutions contained (in mM): 140 NaCl, 4 KCl, 10 HEPES, 10 D-Glucose, 1 MgCl_2_, 1.5 CaCl_2_, pH 7.4 with NaOH. Data were analyzed using Origin (OriginLab) and GraphPad Prism (GraphPad Software).

The voltage dependence of ion current (I-V) was analyzed using a protocol consisting of steps from a holding potential of -120 mV (for 100 ms) to voltages ranging from -90 to +80 mV for 50 ms in 5 mV increment. The linear component of leaky currents and capacitive transients were subtracted using the P/4 procedure. In the activation and conductance density calculation, we used the equation, G= I/(V-Vr), where Vr (the reversal potential) represents the voltage at which the current is zero. For the activation curves, conductance (G) was normalized and plotted against the voltage from -90 mV to +20 mV. To obtain the conductance density curves, G was divided by the capacitance (C), and plotted against the voltage from -90 mV to +20 mV. For voltage dependence of inactivation, cells were clamped at a holding potential of -90 mV, and were applied to step pre-pulses from -120 mV to +20 mV for 100 ms with an increment of 5 mV. Then, the Na^+^ current was recorded at the test pulse of 0 mV for 50 ms. The peak currents under the test pulses were normalized and plotted against the pre-pulse voltage. Activation and inactivation curves were fit to a Boltzmann function to obtain V_1/2_ and slope values. Time course of inactivation data from the peak current at 0 mV was fitted to a single exponential equation: y = A1 exp(−x/τ_inac_) + y0, where A1 was the relative fraction of current inactivation, τ_inac_ was the time constant, x was the time, and y0 was the amplitude of the steady-state component. Late sodium current was measured as the mean inward current between 40-50 ms at the end of a 100-ms depolarization to voltage between -30 mV and +30 mV. Then the current was divided by the peak inward current at the same potential to show late sodium current percentage.

All data points are presented as mean ± standard error of the mean (SEM) and *n* is the number of experimental cells from which recordings were obtained. Statistical significance was assessed using an unpaired t-test with Welch’s correction, two-way ANOVA analysis and extra sum-of-squares F test.

### Cryo-EM Data Acquisition

For the cryo-EM sample preparation and data collection, the same machine and software were used with similar protocols as previously described (37). Aliquots of 3.5 μL freshly purified Na_v_1.5-E1784K were placed on glow-discharged holey carbon grids (Quantifoil Au 300 mesh, R1.2/1.3). Grids were blotted for 3.0 s and plunge frozen in liquid ethane cooled by liquid nitrogen with Vitrobot Mark IV (Thermo Fisher). Electron micrographs were acquired on a Titan Krios electron microscope (Thermo Fisher) operating at 300 kV, Gatan K3 Summit detector and GIF Quantum energy filter. A total of 4,849 movie stacks were automatically collected, using AutoEMation (46) with a slit width of 20 eV on the energy filter and a preset defocus range from -1.8 μm to -1.3 μm in super-resolution mode at a nominal magnification of 81,000 ×. Each stack was exposed for 2.56 s with 0.08 s per frame, resulting in 32 frames per stack. The total dose rate was 50 e^-^/Å^2^ for each stack. The stacks were motion corrected with MotionCor2 (47) and binned 2-fold, resulting in 1.0825 Å/pixel. Meanwhile, dose weighting was performed (48). The defocus values were estimated with Gctf (49).

### Image Processing

A diagram for the workflow of data processing is presented in *SI Appendix* Fig. S2. A total of 1,471,342 particles were automatically picked from 4,633 manually selected micrographs using Gautomatch. And subsequent two-dimension (2D) and three-dimension (3D) classifications and refinement were performed using Cryosparc (50). After 2D classification, 276,241 good particles were selected and applied to 3D homogeneous refinement. After that one additional round of heterogeneous refinement was performed, during which those particles were classified into 4 classes and the good class was selected, resulting in a dataset of 147,600 particles. The selected particles were applied to the uniform refinement and non-uniform refinement procedure, and finally yielding a 3D EM map with an overall resolution of 3.3 Å. Resolution was estimated with the gold-standard Fourier shell correlation 0.143 criterion (51) with high resolution noise substitution (52).

### Model Building and Structure Refinement

The initial model of Na_v_1.5-E1784K was based on the coordinate of human Na_v_1.5-quinidine, and the structure refinement process was the same as that used before (16, 37) (PDB accession number: 6LQA), was fitted into the EM map by CHIMERA (53) and manually adjusted in COOT (54). The chemical properties of amino acids were taken into consideration during model building. There was no density corresponding to β1. In total, 1,151 residues in Na_v_1.5-E1784K were assigned with side chains, 9 sugar moieties were built. The N-terminal 118 residues, intracellular I-II linker (residues 430-698), II-III linker (residues 945-1187), and C-terminal sequences after Ser1782 were not modelled due to the lack of corresponding densities.

Structure refinement was performed using phenix.real_space_refine application in PHENIX (55) real space with secondary structure and geometry restraints. Over-fitting of the overall model was monitored by refining the model in one of the two independent maps from the gold-standard refinement approach and testing the refined model against the other map (56). Statistics of the map reconstruction and model refinement can be found in *SI Appendix* Table S5.

## Acknowledgements

We thank Xiaomin Li (Tsinghua University) for technical support during EM image acquisition. This work was funded by the National Key R&D Program (2016YFA0500402 to X.P.) from Ministry of Science and Technology of China, and Beijing Nova Program (Z191100001119127) from Beijing Municipal Science and Technology Commission. We thank the Tsinghua University Branch of China National Center for Protein Sciences (Beijing) for providing the cryo-EM facility support. We thank the computational facility support on the cluster of Bio-Computing Platform (Tsinghua University Branch of China National Center for Protein Sciences Beijing) and the “Explorer 100” cluster system of Tsinghua National Laboratory for Information Science and Technology. N.Y. is supported by the Shirley M. Tilghman endowed professorship from Princeton University.

## Author contributions

N.Y. conceived the project. Z.L., X.Z., W.W., X.P., and J.L. performed experiments for structural determination. X.J., T.W., and X.P. performed and analyzed electrophysiological measurements. All authors contributed to data analysis. N.Y. wrote the manuscript.

## Notes

### Competing Interest Statement

The authors have declared no competing interest.

