## Supplemental information for "Structure of human Na_v_1.5 reveals the fast inactivation-related segments as a mutational hotspot for the Long QT Syndrome"

### Supplementary Information

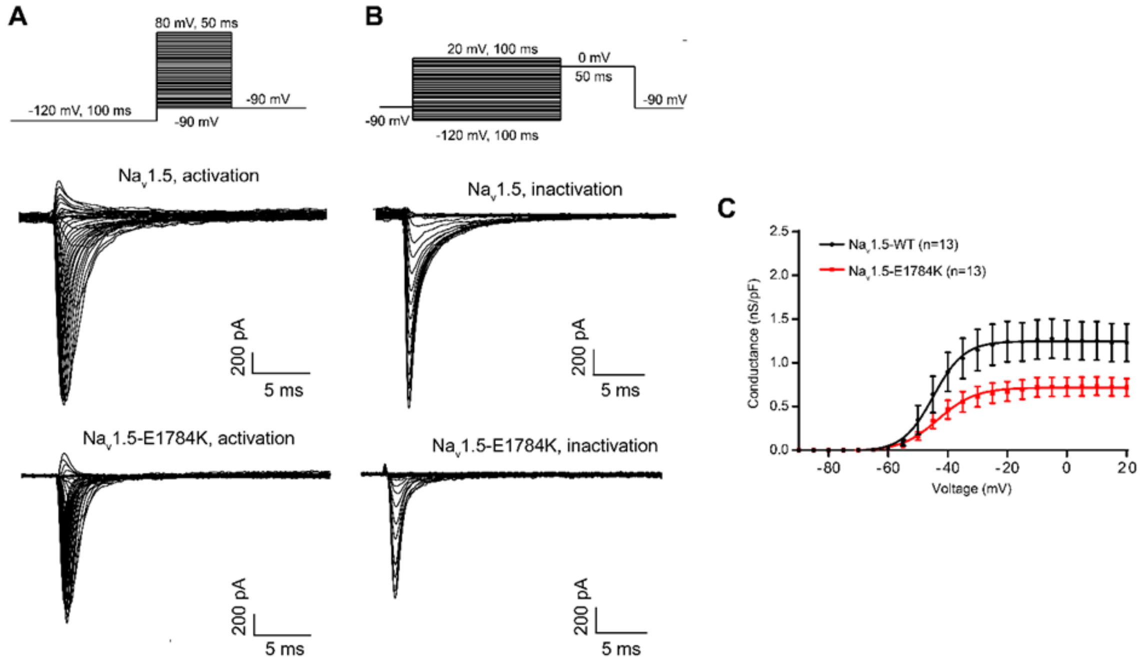


**Figure S1 | Electrophysiological properties of human Na$_V$1.5 and Na$_V$1.5-E1784K transiently expressed in HEK293T cells.** (*A)* Representative whole-cell voltage-dependent activation traces obtained from HEK293T cells transfected with either human Na$_V$1.5 or Na$_V$1.5-E1784K. (*B*) Voltage-dependent inactivation traces. The top panels show the diagrams of recording protocols in (*A*) and (*B*). (*C*) Conductance density. Na$_V$1.5-E1784K significantly decreased $Na^+$ current compared with Na$_V$1.5. *n* values indicate the number of independent cells recorded; mean ± s.e.m. Detailed parameters are presented in Table S4.

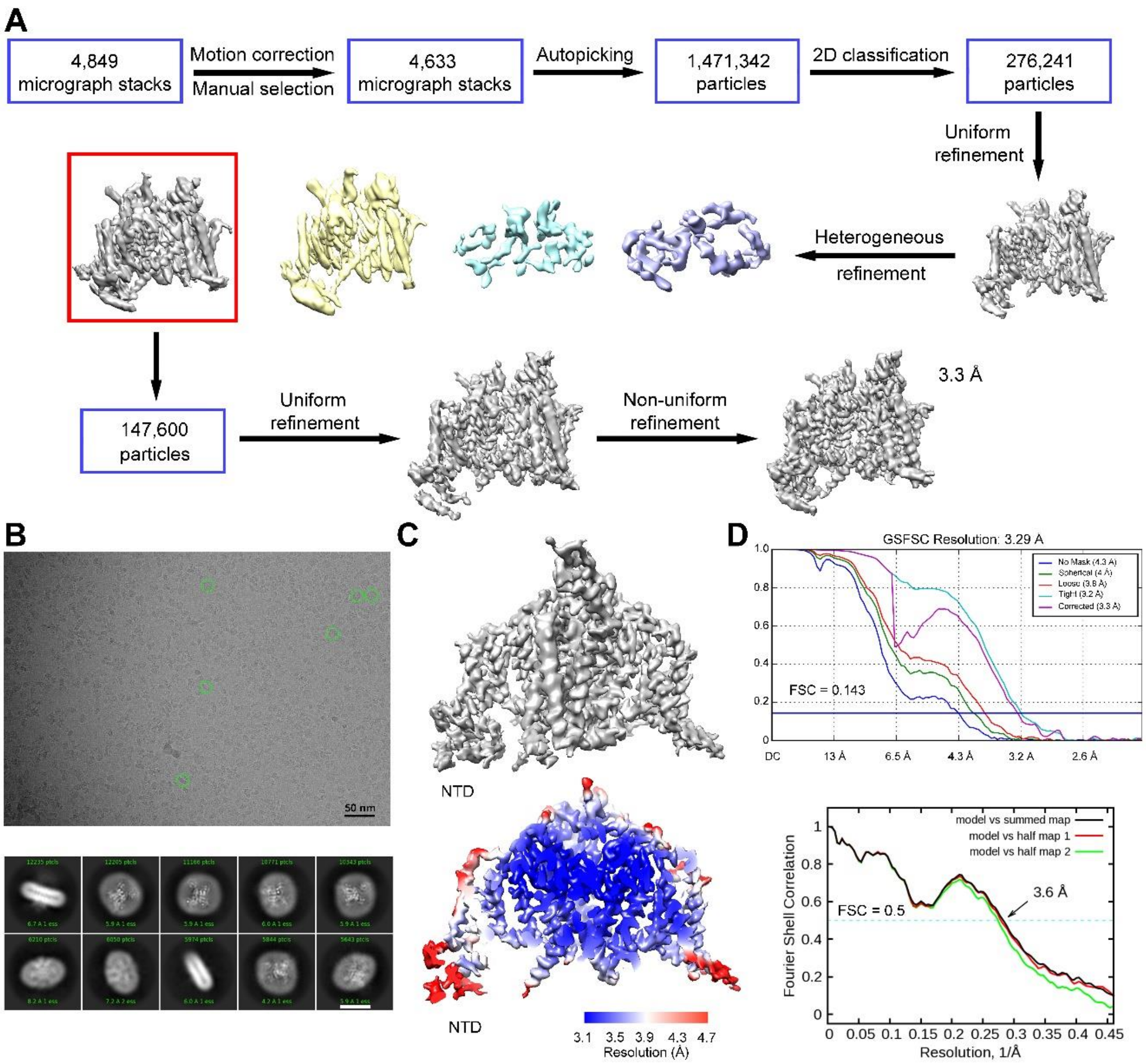


**Figure S2 | Cryo-EM analysis of human Na$_v$1.5-E1784K.** (*A*) The flowchart for EM data processing. Details can be found in Materials and Methods. (*B*) Representative electron micrograph and two-dimensional class averages of Na$_v$1.5-E1784K. Particles in distinct orientations are highlighted by green circles. Scale bar represents 50 nm. White scale bars represent 10 nm in the lower pannel. (*C*) The 3D EM reconstructions of Na$_v$1.5-E1784K. The maps were generated in CHIMERA. *(D)* Gold-standard Fourier shell correlation (FSC) curve for the 3D reconstruction of Na$_v$1.5-E1784K. FSC curves of the refined model versus the overall map that was refined against (black), of the model refined in the first of the two independent maps used for the gold-standard FSC versus that same map (red), and of the model refined in the first of the two independent maps versus the second independent map (green). The small difference between the red and green curves indicates that the refinement of the atomic coordinates did not suffer from overfitting.

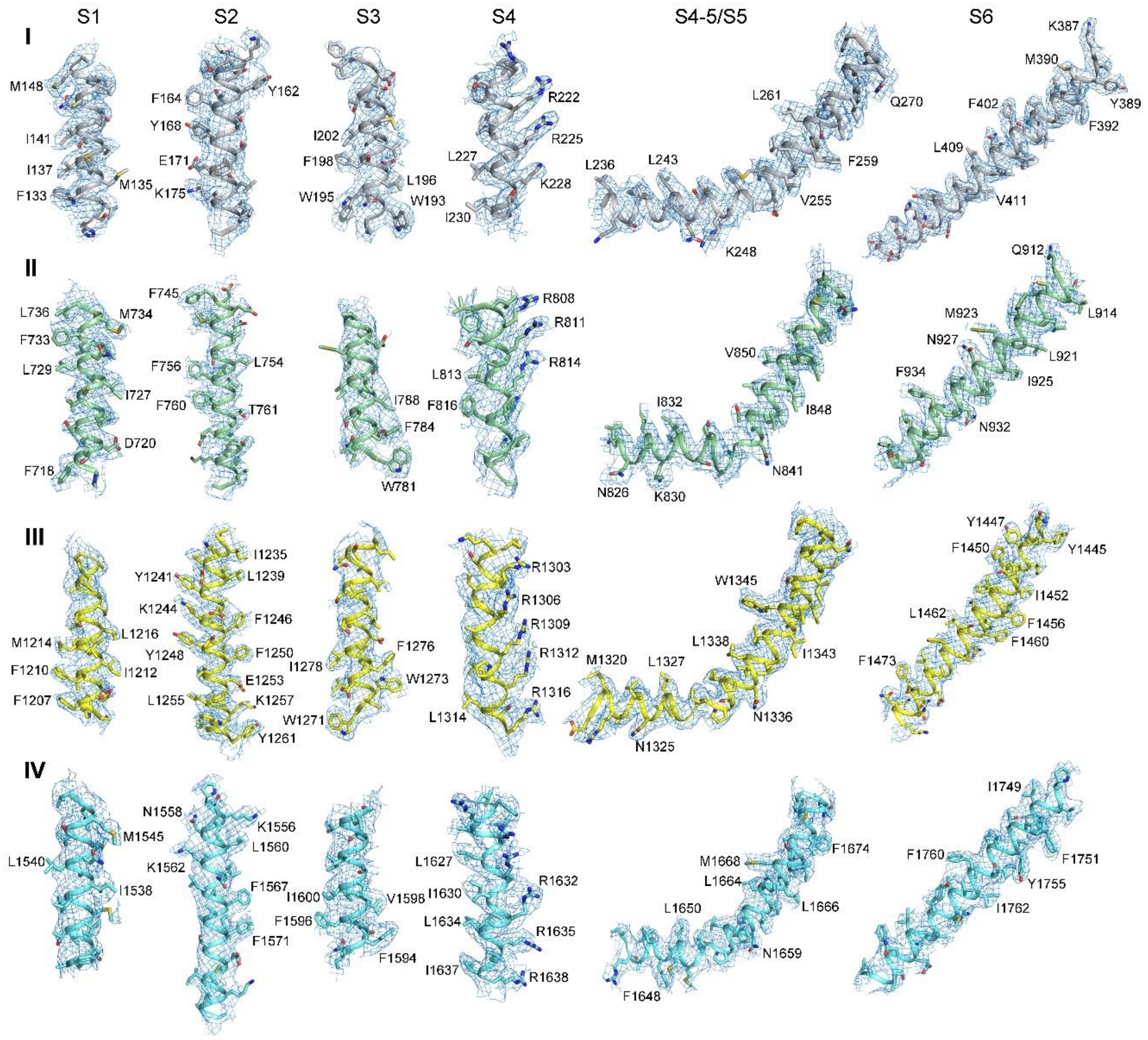


**Figure S3 | EM maps for representative segments of $Na_v1.5$-E1784K.**

EM maps for the S1-S6 segments in each repeat of $Na_v1.5$-E1784K. The maps were prepared in Pymol and contoured at 4 σ.

**Table S1 | Mutations on human $Na_V1.5$ that are associated with Long QT syndrome type 3 (LQT 3).**

| Mutations | Structure | Mutations | Structure | Mutations | Structure |
|---|---|---|---|---|---|
| G9V | NTD | A572V | I-II linker | T1304M | S4III |
| R18Q | NTD | Q573E | I-II linker | N1325S | S4-5III |
| R27H | NTD | G579R | I-II linker | A1326S | S4-5III |
| E30G | NTD | G615E | I-II linker | A1330P | S4-5III |
| R43Q | NTD | L619F | I-II linker | A1330T | S4-5III |
| E48K | NTD | P637L | I-II linker | P1332L | S5III |
| P52S | NTD | G639R | I-II linker | S1333Y | S5III |
| R53Q | NTD | P648L | I-II linker | I1334V | S5III |
| R104G | NTD | E654K | I-II linker | L1338V | S5III |
| S115G | NTD | L673P | I-II linker | R1432S | L6III |
| V125L | NTD | R680H | I-II linker | S1458Y | S6III |
| L212P | S3-4I | R689C | I-II linker | N1472S | S6III |
| R222Q | S4I | R689H | I-II linker | F1473C | S6III |
| R225Q | S4I | P701L | I-II linker | G1481E | III-IV linker |
| R225W | S4I | T731I | S1II | F1486L | III-IV linker |
| V240M | S4-5I | Q750R | S2II | M1487L | III-IV linker |
| Q245K | S4-5I | D772N | S2-3II | T1488R | III-IV linker |
| V247L | S4-5I | F816Y | S4II | E1489D | III-IV linker |
| N275K | L5I | I848F | S5II | K1493R | III-IV linker |
| G289S | L5I | S941N | S6II | Y1495S | III-IV linker |
| R340W | L5I | Q960K | II-III linker | M1498V | III-IV linker |
| R367C | P1I | R965L | II-III linker | L1501V | III-IV linker |
| T370M | P1I | R971C | II-III linker | K1505N | III-IV linker |
| I397T | S6I | C981F | II-III linker | V1532I | S1IV |
| L404Q | S6I | A997S | II-III linker | L1560F | S2IV |
| N406K | S6I | C1004R | II-III linker | I1593M | S3IV |
| L409V | S6I | E1053K | II-III linker | F1594S | S3IV |
| V411M | S6I | T1069M | II-III linker | F1596I | S3IV |
| A413E | S6I | A1100V | II-III linker | S1609W | S3IV |
| A413T | S6I | D1114N | II-III linker | T1620K | S4IV |
| E462A | I-II linker | D1166N | II-III linker | R1623L | S4IV |
| E462K | I-II linker | R1193Q | II-III linker | R1623Q | S4IV |
| F530V | I-II linker | Y1199S | II-III linker | R1626H | S4IV |
| R535Q | I-II linker | E1225K | S1-2III | R1626P | S4IV |
| R569W | I-II linker | E1231K | S1-2III | R1644C | S4-5IV |
| S571I | I-II linker | F1250L | S2III | R1644H | S4-5IV |
| A572D | I-II linker | L1283M | S3III | T1645M | S4-5IV |
| A572S | I-II linker | E1295K | S3-4III | L1650F | S4-5IV |

| Mutations | Structure | Mutations | Structure | Mutations | Structure |
|---|---|---|---|---|---|
| M1652R | S4-5IV | E1784K | CTD | R1913H | CTD |
| M1652T | S4-5IV | D1790G | CTD | A1949S | CTD |
| I1660V | S5IV | Y1795C | CTD | V1951L | CTD |
| V1667I | S5IV | Y1795YD | CTD | Y1977N | CTD |
| T1723N | L6IV | D1819N | CTD | F2004L | CTD |
| R1739W | L6IV | L1825P | CTD | F2004V | CTD |
| L1761F | S6IV | R1826H | CTD | R2012C | CTD |
| L1761H | S6IV | D1839G | CTD | ΔA586-L587 | I-II linker |
| V1763M | S6IV | H1849R | CTD | ΔE429 | S6I |
| M1766L | S6IV | R1897W | CTD | ΔI1212 | S1III |
| Y1767C | S6IV | E1901Q | CTD | ΔK1505-Q1507 | III-IV linker |
| I1768V | S6IV | S1904L | CTD | ΔQ1507-P1509 | III-IV linker |
| V1777M | S6IV | Q1909R | CTD | ΔF1617 | S3-4IV |
| T1779M | S6IV | | | | |

Disease mutations that structurally unresolved regions are shaded light gray. The same applies to Tables S2 and S3.

Mutations in Tables S1-S3 are summarized from https://www.uniprot.org/uniprot/Q14524.

**Table S2 | Mutations on human $Na_V1.5$ that are associated with Brugada syndrome.**

| Mutations | Structure |
|---|---|
| R18Q | NTD |
| R27H | NTD |
| N70K | NTD |
| D84N | NTD |
| F93S | NTD |
| I94S | NTD |
| V95I | NTD |
| R104Q | NTD |
| R104W | NTD |
| N109K | NTD |
| R121Q | NTD |
| R121W | NTD |
| K126E | NTD |
| L136P | S1I |
| V146M | S1I |
| E161K | S2I |
| E161Q | S2I |
| K175N | S2I |
| A178G | S2I |
| C182R | S2-3I |
| A185V | S2-3I |
| T187I | S2-3I |
| A204V | S3I |
| L212Q | S3-4I |
| T220I | S4I |
| R222Q | S4I |
| V223L | S4I |
| A226V | S4I |
| I230T | S4I |
| V232I | S4-5I |
| V240M | S4-5I |
| Q270K | S5I |
| L276Q | L5I |
| H278D | L5I |
| R282C | L5I |
| R282H | L5I |
| V294M | L5I |
| V300I | L5I |
| L315P | L5I |
| G319S | L5I |
| T320N | L5I |
| L325R | L5I |
| P336L | L5I |
| G351D | L5I |
| G351V | L5I |
| T353I | L5I |
| D356N | L5I |
| R367C | P1I |
| R367H | P1I |
| R367L | P1I |
| M369K | P1I |
| W374G | P2I |
| R376H | P2I |
| G386E | L6I |
| G386R | L6I |
| V396A | S6I |
| V396L | S6I |
| N406S | S6I |
| E439K | I-II linker |
| D501G | I-II linker |
| G514C | I-II linker |
| R526H | I-II linker |
| F532C | I-II linker |
| F543L | I-II linker |
| G552R | I-II linker |
| L567Q | I-II linker |
| G615E | I-II linker |
| L619F | I-II linker |
| R620C | I-II linker |
| T632M | I-II linker |
| P640A | I-II linker |
| A647D | I-II linker |
| P648L | I-II linker |
| R661W | I-II linker |
| H681P | I-II linker |
| C683G | I-II linker |
| P701L | I-II linker |
| P717L | I-II linker |
| A735E | S1II |
| A735V | S1II |
| E746K | S1-2II |
| G752R | S2II |
| G758E | S2II |
| M764R | S2II |
| D772N | S2-3II |
| P773S | S2-3II |
| V789I | S3II |
| R808P | S4II |
| L812Q | S4II |
| R814Q | S4II |
| K817E | S4II |
| L839P | S5II |
| F851L | S5II |
| E867Q | L5II |
| R878C | L5II |
| R878H | L5II |
| H886P | P1II |
| F892I | P1II |
| R893C | P1II |
| R893H | P1II |
| C896S | P1II |
| E901K | P2II |
| S910L | L6II |
| C915R | S6II |
| L917R | S6II |
| N927S | S6II |
| L928P | S6II |
| L935P | S6II |
| R965C | II-III linker |
| R965H | II-III linker |
| A997T | II-III linker |
| R1023H | II-III linker |
| E1053K | II-III linker |
| D1055G | II-III linker |

| Mutations | Structure |
|---|---|
| S1079Y | II-III linker |
| A1113V | II-III linker |
| S1140T | II-III linker |
| R1193Q | II-III linker |
| S1219N | S1III |
| E1225K | S1-2III |
| Y1228H | S1-2III |
| R1232Q | S1-2III |
| R1232W | S1-2III |
| K1236N | S2III |
| L1239P | S2III |
| D1243N | S2III |
| V1249D | S2III |
| E1253G | S2III |
| G1262S | S2-3III |
| W1271C | S3III |
| A1288G | S3III |
| F1293S | S3-4III |
| L1311P | S4III |
| G1319V | S4-5III |
| V1323G | S4-5III |
| P1332L | S5III |
| F1344L | S5III |
| F1344S | S5III |
| L1346S | S5III |
| L1346P | S5III |
| M1351R | S5III |
| V1353M | S5III |
| G1358W | L5III |
| K1359N | L5III |
| F1360C | L5III |
| C1363Y | L5III |
| S1382I | L5III |
| V1405L | P1III |
| V1405M | P1III |
| G1406E | P1III |
| G1406R | P1III |
| G1408R | P1III |
| Y1409C | P1III |

| Mutations | Structure |
|---|---|
| L1412F | P1III |
| K1419E | P1III |
| G1420R | SF III |
| A1427S | P2III |
| A1428V | P2III |
| R1432G | L6III |
| R1432S | L6III |
| G1433V | L6III |
| P1438L | L6III |
| E1441Q | L6III |
| I1448L | S6III |
| I1448T | S6III |
| Y1449C | S6III |
| V1451D | S6III |
| N1463Y | S6III |
| V1468F | S6III |
| Y1494N | III-IV linker |
| L1501V | III-IV linker |
| G1502S | III-IV linker |
| R1512W | III-IV linker |
| I1521K | III-IV linker |
| V1525M | III-IV linker |
| K1527R | III-IV linker |
| E1548K | S1-2IV |
| A1569P | S2IV |
| F1571C | S2IV |
| E1574K | S2IV |
| L1582P | S2-3IV |
| R1583C | S2-3IV |
| R1583H | S2-3IV |
| V1604M | S3IV |
| Q1613L | S3-4IV |
| T1620M | S3-4IV |
| R1623Q | S4IV |
| R1629Q | S4IV |
| G1642E | S4-5IV |
| R1644C | S4-5IV |
| A1649V | S4-5IV |
| I1660V | S5IV |

| Mutations | Structure |
|---|---|
| G1661R | S5IV |
| V1667I | S5IV |
| S1672Y | S5IV |
| A1680T | L5IV |
| D1690N | L5IV |
| A1698T | P1IV |
| T1709M | P1IV |
| T1709R | P1IV |
| G1712S | SFIV |
| D1714G | P2IV |
| N1722D | L6IV |
| C1728R | L6IV |
| C1728W | L6IV |
| G1740R | L6IV |
| G1743E | L6IV |
| G1743R | L6IV |
| G1748D | S6IV |
| V1764F | S6IV |
| T1779M | S6IV |
| E1784K | CTD |
| Y1795H | CTD |
| Y1795YD | CTD |
| Q1832E | CTD |
| C1850S | CTD |
| V1861I | CTD |
| K1872N | CTD |
| V1903L | CTD |
| A1924T | CTD |
| G1935S | CTD |
| E1938K | CTD |
| V1951L | CTD |
| I1968S | CTD |
| F2004L | CTD |
| F2004V | CTD |
| ΔF393 | S6I |
| ΔK1479 | III-IV linker |
| ΔK1500 | III-IV linker |
| ΔF1617 | S3-4IV |

**Table S3 | Mutations on human $Na_V1.5$ that are associated with cardiac disorders other than LQT3 and Brugada syndrome.**

| Mutations | Disease | Structure |
|---|---|---|
| E161K | PFHB1A | S2I |
| R225W | PFHB1A | S4I |
| G298S | PFHB1A | L5I |
| T512I | PFHB1A | I-II linker |
| G514C | PFHB1A | I-II linker |
| G752R | PFHB1A | S2II |
| R1232W | PFHB1A | S1-2III |
| D1595N | PFHB1A | S3IV |
| T1620K | PFHB1A | S4IV |
| T220I | SSS1 | S4I |
| A735V | SSS1 | S1II |
| P1298L | SSS1 | S3-4III |
| G1408R | SSS1 | P1III |
| D1792N | SSS1 | CTD |
| S1710L | VF1 | SFIV |
| F532C | SIDS | I-II linker |
| G1084S | SIDS | II-III linker |
| S1333Y | SIDS | S4-5III |
| F1705S | SIDS | P1IV |
| D1275N | ATRST1 | S3III |
| D1275N | CMD1E | S3III |
| M138I | ATFB10 | S1I |
| E428K | ATFB10 | S6I |
| H445D | ATFB10 | I-II linker |
| N470K | ATFB10 | I-II linker |
| A572D | ATFB10 | I-II linker |
| E655K | ATFB10 | I-II linker |
| T1131I | ATFB10 | II-III linker |
| R1826C | ATFB10 | CTD |
| V1951M | ATFB10 | CTD |
| N1987K | ATFB10 | CTD |

**PFHB1A**: Progressive familial heart block 1A; **SSS1**: Sick sinus syndrome; **VF1**: Familial paroxysmal ventricular fibrillation 1; **SIDS**: Sudden infant death syndrome; **ATRST1**: Atrial standstill 1; **CMD1E**: Cardiomyopathy, dilated 1E; **ATFB10**: Atrial fibrillation, familial, 10.

**Table S4 | Activation and steady-state inactivation parameters of $Na_v1.5$ and $Na_v1.5$-E1784K in HEK293T cells.**

| | Parameters | $Na_v1.5$-WT | $Na_v1.5$- E1784K |
|---|---|---|---|
| **Activation** | $V_{1/2}$ (mV) | -41.20 ± 0.37 | -35.88 ± 0.54$^{****}$ |
| | P | / | < 0.0001 |
| | slope | 6.44 ± 0.32 | 9.37 ± 0.48$^{****}$ |
| | P | / | < 0.0001 |
| | n | 15 | 17 |
| **Inactivation** | $V_{1/2}$ (mV) | -77.28 ± 0.73 | -96.07 ± 0.66**** |
| | P | / | < 0.0001 |
| | slope | -12.77 ± 0.66 | -9.91 ± 0.46*** |
| | P | / | 0.0003 |
| | $\tau_{inac}$ (ms) | 1.47 ± 0.18 | 0.78 ± 0.05*** |
| | P | / | 0.0009 |
| | n | 12 | 13 |
| **Conductance** | $G_{top}$ (nS/pF) | 1.25 ± 0.06 | 0.72 ± 0.03*** |
| | P | / | 0.0004 |
| | n | 13 | 13 |
| **Late sodium current** | Late current (%) | 0.71 ± 0.15 | 1.84 ± 0.29* |
| | P | / | 0.0425 |
| | n | 10 | 12 |

* P < 0.05 versus WT, ** P < 0.01 versus WT, *** P < 0.001 versus WT, **** P < 0.0001 versus WT. Each data point represents mean ± s.e.m (standard deviation of mean) and *n* is the number of experimental cells from which recordings were obtained. The extra sum-of-squares F test was used to compare the $V_{1/2}$ of activation and inactivation fits and $G_{top}$ of conductance fits. $\tau_{inac}$ values of $Na_v1.5$ and $Na_v1.5$-E1784K were compared using an unpaired t-test with Welch's correction. Late sodium current (%) was compared using two-way ANOVA analysis at voltage from -30 mV to 30 mV (Late current percentage at -30 mV was shown in the table).

**Table S5 | Cryo-EM data collection, refinement and validation statistics**

| | |
|---|---|
| **Data collection** | |
| EM equipment | Titan Krios (Thermo Fisher) |
| Voltage (kV) | 300 |
| Detector | Gatan K3 Summit |
| Energy filter | Gatan GIF Quantum, 20 eV slit |
| Pixel size (Å) | 1.0825 |
| Electron dose ($e^-$/Å$^2$) | 50 |
| Defocus range (μm) | 1.3 ~ 1.8 |
| Number of collected micrographs | 4,849 |
| Number of used micrographs | 4,633 |
| **Reconstruction** | |
| Software | Cryosparc |
| Number of used particles | 147,600 |
| Symmetry | C1 |
| Resolution (Å) | 3.3 |
| Map sharpening B-factor (Å$^2$) | -88.3 |
| **Refinement** | |
| Software | Phenix |
| Cell dimensions | |
| a=b=c (Å) | 259.80 |
| α=β=γ (˚) | 90 |
| Model composition | |
| Protein residues | 1,151 |
| Side chains assigned | 1,151 |
| Sugars | 9 |
| R.m.s deviations | |
| Bonds length (Å) | 0.003 |
| Bonds Angle (˚) | 0.621 |
| Ramachandran plot statistics (%) | |
| Favored | 91.44 |
| Allowed | 8.12 |
| Outlier | 0.44 |
